# Suspension physics govern the multiscale dynamics of blood flow in sickle cell disease

**DOI:** 10.1101/2025.03.13.642599

**Authors:** Hannah M. Szafraniec, Freya Bull, John M. Higgins, Howard A. Stone, Timm Krüger, Philip Pearce, David K. Wood

## Abstract

In diseases from diabetes to malaria, blood dynamics are significantly altered, resulting in poor clinical outcomes. However, the multiscale mechanisms that determine blood flow in the microcirculation in health and disease are undefined, largely owing to the difficulty in directly linking cell properties to whole-blood rheology. Here, we overcome these difficulties by developing a microfluidic platform to measure red blood cell properties and flow dynamics in the same blood samples from donors. We focus on sickle cell disease (SCD), a genetic disorder that causes red blood cells to stiffen in deoxygenated conditions, with disease pathology driven by oxygen-dependent blood rheology. Our linked cell and whole-blood measurements establish that increases in effective resistances in heterogeneous suspensions are driven by increases in the proportion of stiff cells, similar macroscopically to the behavior of rigid-particle suspensions. Furthermore, by combining simulations with spatially resolved measurements of cell dynamics, we show how the spatio-temporal organization of stiff and deformable cells determines blood rheology and drives disease pathophysiology. In the presence of deformable cells, the stiffened cells marginate towards channel walls, increasing effective wall friction. In fully deoxygenated conditions in which all cells are stiffened, significant heterogeneity in cell volume fraction along the direction of flow causes localized jamming, drastically increasing effective viscous flow resistance. Our work defines the relevant suspension physics required to understand pathological blood rheology in SCD and other diseases affecting red blood cell properties. More broadly, we reveal the multiscale processes that determine emergent rheology in heterogeneous particle suspensions.

---

A common feature of cancer^1^, metabolic diseases such as diabetes^2^, and infectious diseases such as malaria^3^ are pathological changes to the properties of blood, which can disrupt nutrient transport and oxygen transport throughout the body. Sickle cell disease (SCD), which affects millions of people worldwide and 300,000 infants born each year, is a prevalent example of a genetic blood disorder^4^. The pathology of SCD derives from a mutation in the oxygen-carrying hemoglobin molecule^5^ causing it to aggregate into polymer under deoxygenation, so that the red blood cells (RBCs) become significantly less deformable^6–10^. The result is that patients with SCD exhibit pathological and variable blood flow, with disease progression and severity linked to rheological and hemodynamic biomarkers, such as elevated viscosity and increased likelihood of vaso-occlusive crises^11–18^. Despite the prevalence of such blood disorders, we still lack a mechanistic understanding of how alterations in the properties or distribution of blood cells alter hemodynamics. For example, it remains unclear why individuals with SCD exhibit significantly variable blood rheology, which is linked to certain clinical outcomes. We also do not understand how the presence of stiff RBCs impacts blood flow in the microvasculature, where most of the hemodynamically driven disease processes, such as vaso-occlusive crisis, are thought to occur. By contrast, extensive work understanding healthy blood flow across multiple scales has aided progress in drug delivery, medical device designs, and clinical risk assessment in diseased vascular structures, such as aneurysms^19–21^. Similar progress in blood disorders is hampered by a lack of experimental measurements and theoretical models that directly connect disease-driven heterogeneity in RBC properties to emergent blood flow dynamics and effective rheology under physiologic conditions.

For SCD, multiple studies using macroscale measurements with rheometers or simulations have shown that increasing the volume fraction of stiffened RBCs increases effective blood viscosity^22–24^. These measurements provide a clear link between stiffened RBCs and increased blood viscosity, but they are difficult to interpret mechanistically and extrapolate to physiologic hemodynamics. Specifically, measurements using rheometers are not spatially resolved, and do not capture the microscopic dynamics of the cells, such as the well-known Fahraeus-Lindqvist and Fahraeus effects which arise from interactions between cells and confining walls^25,26^. How such phenomena are affected by heterogeneous RBC properties is not captured by macroscale measurements of effective viscosity. Although models have been developed that aim to accurately predict blood flow in confinement, they typically do so with the assumption that all cells are homogeneous and highly deformable^27–31^. More generally, suspensions of homogeneous rigid particles or cells have been well-characterized experimentally, theoretically, and computationally^32–36^. These studies have revealed that effects arising from the suspended particles themselves are significant and, for example, lead to radial variations in particle concentration that cause apparent non-Newtonian rheological behavior in confined flow^37,38^. Thus, both experiments and modeling in blood and suspensions of rigid particles have revealed that microscopic effects, owing to the red blood cells and rigid particles, respectively, contribute significantly to suspension rheology and flow dynamics. Therefore, we should expect heterogeneous mixtures of RBCs to have unique microscopic effects, but these phenomena are not well-represented in existing models. As a result, we do not understand how sickle blood flows in physiologically relevant conditions, including the microvasculature, where complications are likely to occur.

In this study, we significantly advance our understanding of SCD hemodynamics under physiologic conditions by mechanistically linking the distribution of RBC mechanical properties with spatially resolved blood flow, both transverse and along the flow directions, in confined geometries. We use a high-throughput imaging method to measure distinct subpopulations of RBCs within individual patient blood samples^39,40^. In parallel, we measure effective rheology of the blood derived from spatially resolved flow fields^41^. To explore mechanisms leading to the emergent rheology, we develop a computational model to simulate confined flow of heterogeneous mixtures of cells. This allows us to probe the dynamics of the cells and measure the impact on rheology, which we confirm experimentally. We are thereby able to directly link single cell properties to blood dynamics in microchannels across individual donors, and to establish mechanisms for the apparent rheology of heterogeneous mixtures. Our findings reveal how suspension physics govern the flow properties of sickle blood and explain differences between patients. Despite the presence of both stiffened and deformable cells in the blood, we find that our measurements are broadly consistent with previous empirical models for the rheology of rigid-particle suspensions across a range of particle volume fractions. Additionally, our imaging and computational modelling reveal how overall dynamics are driven by complex spatio-temporal processes in mixtures of stiff and deformable cells. Therefore, our work elucidates the dominant physical parameters and mechanisms contributing to pathological rheology for patients with SCD. More broadly, our findings advance understanding of the flow of particle suspensions by revealing how rheological properties emerge from heterogeneity in particle properties.

## Stiffened RBCs are the key driver of changes in SCD patient rheology across oxygen tensions and between patients

Using single-cell measurements in parallel with a microfluidic rheological platform, we combined measurements of cell properties – the fraction of RBCs that are stiff due to hemoglobin S polymerization – with whole blood dynamics – cell tracks, effective flow fields, and overall flow resistance (R_effective_, i.e., the ratio of pressure drop to flow rate normalized to the 21% oxygen value) – over a range of experimental oxygen tensions, within the same samples from multiple patients (Fig. 1a, b, Material and Methods 1.0-5.0). We hypothesized that the proportion of stiff RBCs drives the rheological response to hypoxia. Given our blood flow experiments have a fixed volume fraction of RBCs (*Φ = 0*.*25*), our results are scaled to the fraction of RBCs that are stiff (χ*stiff*), where χ_*stiff*_ *= Φ*_*stiff*_ */Φ*. While both R_effective_ and χ_*stiff*_ increased with decreasing oxygen, each patient exhibited a unique response curve (Fig. 1c, d). By plotting R_effective_ against χ_*stiff*_, we found all samples collapsed onto a single curve, including patient rheology from exchange transfusion treatment (Fig. 1e, Materials and Methods 2.0). Therefore, for a fixed volume fraction of RBCs in suspension, these results indicate that the fraction of stiff RBCs is the key microscopic driver of whole-blood rheology across SCD patients and over the full physiologic range of oxygen tensions.

**Figure 1.**
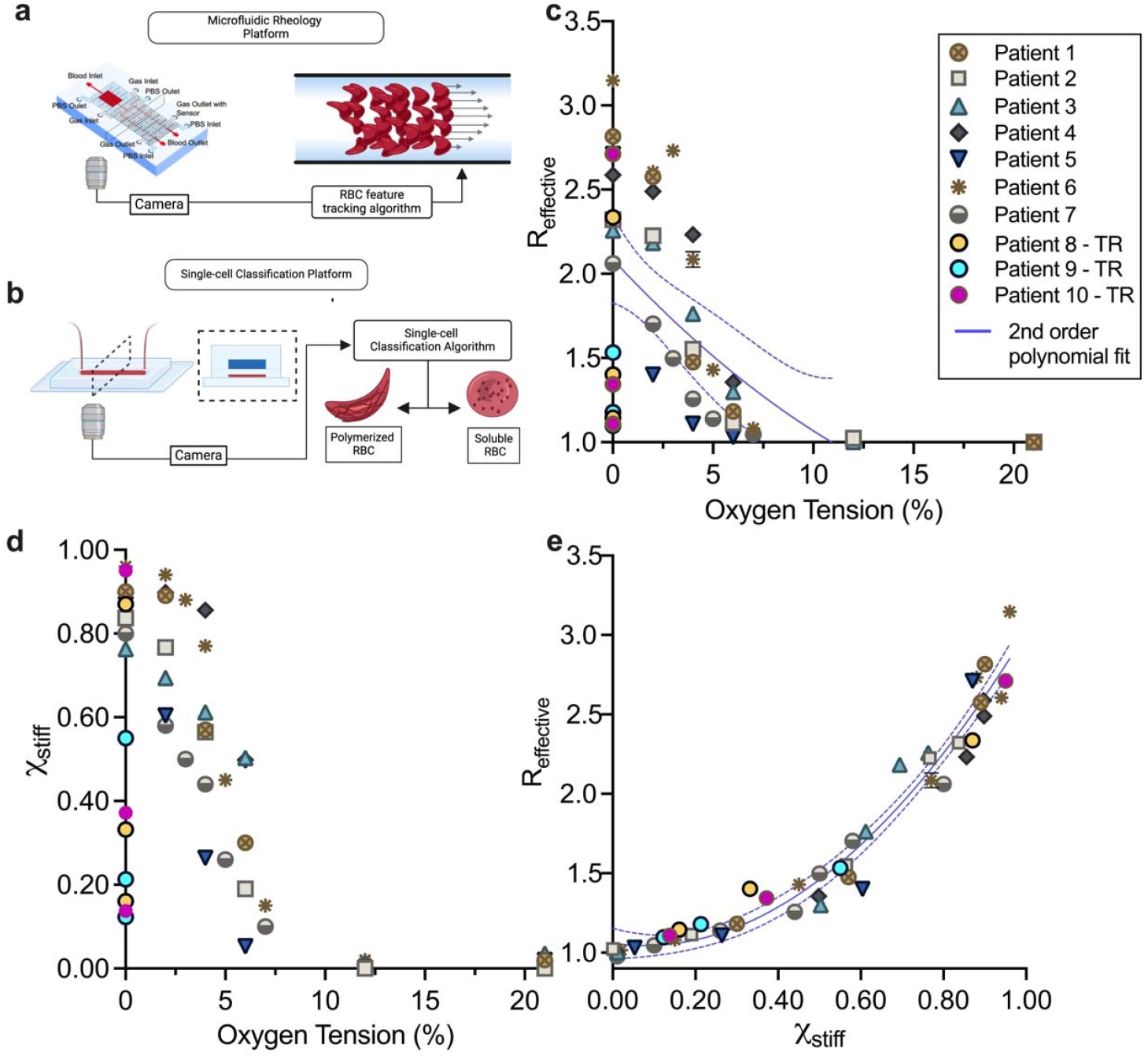
Stiff cells drive the increase in effective resistance for sickle blood under hypoxia and reduce inter-patient variability. a) The microfluidic rheology platform used to measure spatially resolved blood flow dynamics. Blood was perfused in a 20×20 μm cross-sectional channel, and R_effective_ was measured from flow data. b) The microfluidic single-cell classification platform used to measure the RBC population distribution of stiff (cells with polymer) and deformable (cells without polymer) cells at various oxygen tensions. c) Effective resistances were normalized by the 21% oxygen value for each patient and plotted against oxygen tension. Results show unique patient response curves at oxygen tensions less than 12%. Patients 8, 9 and 10 were transfused twice and measurements were performed at 0% oxygen, including the native sample. The variance measured by the sum of square errors of a 2^nd^ order polynomial curve fit was 0.28. d) Stiff cell fractions plotted for each patient across a range of oxygen tensions. e) Effective resistance plotted against stiff cell fraction decreased inter-patient rheological variability by a factor of 17 with a variance of 0.020. Results shown for n = 10 sickle blood samples. Error bars represent plus or minus the standard error of the mean (+/-SEM) for resistance data gathered for n = 11 independent sampling time points during flow data acquisition. Error bars smaller than the data symbol are not shown. Statistical analysis performed in Prism v9.5.0 using nonlinear least-squares regression of a 2^nd^ order polynomial. 2^nd^ order polynomial fits (blue line) are plotted with 95% confidence bands (dashed lines).

## Whole blood from SCD patients behaves macroscopically as a suspension of rigid particles with effective properties

Next, we explored the physical basis for the relationship between stiff cell fraction and effective flow resistance. Our spatially resolved flow measurements allow for the separation of the effective resistance into a frictional element (R_*friction*_), associated with RBCs slipping near channel walls, and a viscous element (R_*viscous*_), which describes the remaining resistance to flow owing to the effective bulk properties of the blood (Fig. 2a, Material and Methods 4.0). Again, we normalize each resistance to the corresponding resistance measured at 21% oxygen. These quantities represent macroscopic measurements of the overall oxygen-dependent effective rheology of the blood. In suspensions of homogeneous rigid particles, a range of experimental measurements have demonstrated that slip and bulk resistances follow established empirical relationships based on the volume fraction of the particles. For suspensions of low aspect ratio particles, the slip length has been measured as a function of particle diameter, particle volume fraction, and the critical particle volume fraction^33^. Here, we apply this relationship in Eq. 1 considering the frictional resistance an equivalent measurement of the inverse slip length and substitute the volume fraction of rigid particles for the directly measured quantity, **χ**_*stiff*_. We also assume the cell diameter to be fixed. The frictional resistance relative to 21% oxygen is thus fit with Eq. 1 using the single parameter, χ_*m,slip*_:

**Figure 2.**
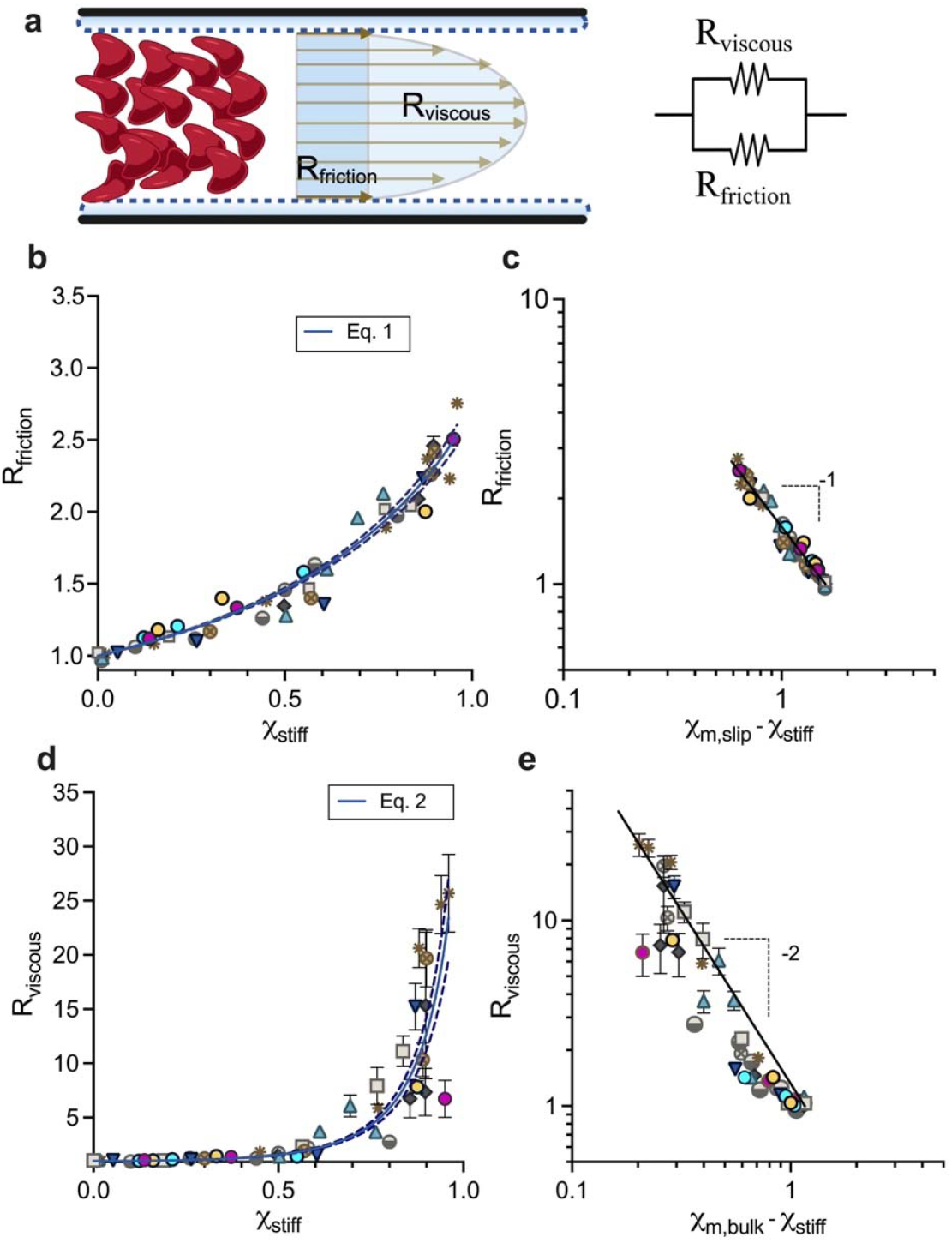
Critical microvascular rheology of SCD blood compared to theory in rigid-particle suspensions. a) Local velocity fields in the microfluidic channel are obtained using high-speed imaging and feature tracking methods to generate flow profiles. Flow profiles approximate the height averaged flow, which is used to measure the effective rheology. With a fixed pressure drop, the frictional resistance, measured from the slip flow rate, and viscous resistance, measured from the bulk flow rate, combine in parallel to form the effective resistance. b) R_*friction*_ plotted against χ _*stiff*_ and fit with Eq.1. c) Logarithmic plot of R_*friction*_ versus (χ _*m,slip*_-χ _*stiff*_) compared to Eq. 1 (black line) shows good qualitative agreement between data and model for a χ _*m,slip*_ of 1.59 (95% CI: 1.56, 1.62). d) R_*viscous*_ versus χ _*stiff*_. Data fit with Eq. 2. e) Logarithmic plot of R_*viscous*_ versus (χ _*m,bulk*_-χ _*stiff*_) compared to Eq. 2 (black line) shows good qualitative agreement between data and model for a χ _*m,bulk*_ of 1.16 (95% CI: 1.15, 1.18). Results shown for n = 10 sickle blood samples. Error bars represent plus or minus the standard error of the mean (+/-SEM) for resistance data gathered for n = 11 independent sampling time points during flow data acquisition. Error bars smaller than the data symbol are not shown. Eq. 1 and Eq. 2 (blue lines) are fit to the data in panel b and d, respectively, using a nonlinear least-squares regression performed in Prism v9.5.0. and plotted with 95% confidence bands (dashed line).

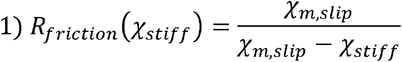

The second feature, viscous resistance, measures the bulk viscosity of the suspension. The bulk viscosity is defined by the volume fraction of particles; however, the relationship is highly dependent on the volume fraction regime. In the dilute limit, the bulk viscosity is linear in the volume fraction^42,43^. As volume fractions increase, the relationship becomes significantly nonlinear and a divergence in the viscosity occurs as the suspension approaches a jamming transition^34^. Here, we apply this relationship in Eq. 2 considering the viscous resistance an equivalent measurement of bulk viscosity, with χ_*stiff*_ substituted for volume fraction, using a single parameter, χ_*m,bulk*:_

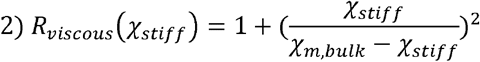

Given the total volume fraction of red blood cells is constant during our experiments, we find the frictional and viscous resistance scale with the fraction of stiff cells in a similar manner to homogenous suspensions with increasing volume fraction of rigid particles. For the frictional resistance, we find the entirety of the data range to be well described by Eq. 1, and we note that increases in the frictional resistance occur for even small amounts of stiff cells (Fig. 2c, d). We also find the viscous resistance to be well described by Eq. 2 (Fig. 2e, f). However, given our estimated blood volume fractions of 25%, the divergence in the viscous resistance seems to occur at a lower volume fraction than predicted from rigid particle suspension theory (S.I. Fig. 1). Overall, our results suggest blood containing stiff RBCs obeys rigid-particle suspension mechanics. However, it is not yet clear what determines the effective parameters describing the critical volume fractions in the well-known empirical relationships.

## In simulations, margination of stiff RBCs increases frictional resistance by increasing the local volume fraction of stiff cells near channel walls

The increases in friction occurring at relatively low stiff RBC fractions suggest extensive interactions between the stiff cells and the channel wall. We hypothesized that complex interactions between deformable and stiff cells could account for this^44^. Therefore, to explore mechanisms for the increases in frictional resistance, we simulated the flow of RBC suspensions composed of mixtures of deformable and stiff cells using a computational model. In our simulations, a lattice-Boltzmann method was used for fluid flow, a finite-element method for cell dynamics, and an immersed boundary method for fluid-structure interactions^30,45,46^. Cells were immersed in a Newtonian fluid and confined to a rectangular channel with flow driven by a constant body force, mimicking a fixed pressure gradient. We performed simulations parameterized from experimental conditions corresponding to a 25% volume fraction of total cells, a 20×20 μm cross-sectional area, and the appropriate pressure drop per unit length (Methods and Materials 7.0). The simulated stiff cell fraction was modulated by increasing the number of stiff cells while decreasing the number of deformable cells. Qualitatively, the simulated suspension demonstrates margination of the stiffer cells toward the channel boundaries (Fig. 3a). Analysis of the local volume fraction of stiff and deformable cells in the simulations show an increase in volume fraction of stiff cells near the channel walls as the stiff cell fraction increases (Fig. 3b-e). Furthermore, by performing resistance analyses on the simulations, we found that this effect drives an increase in friction near the wall (Fig. 3f).

**Figure 3.**
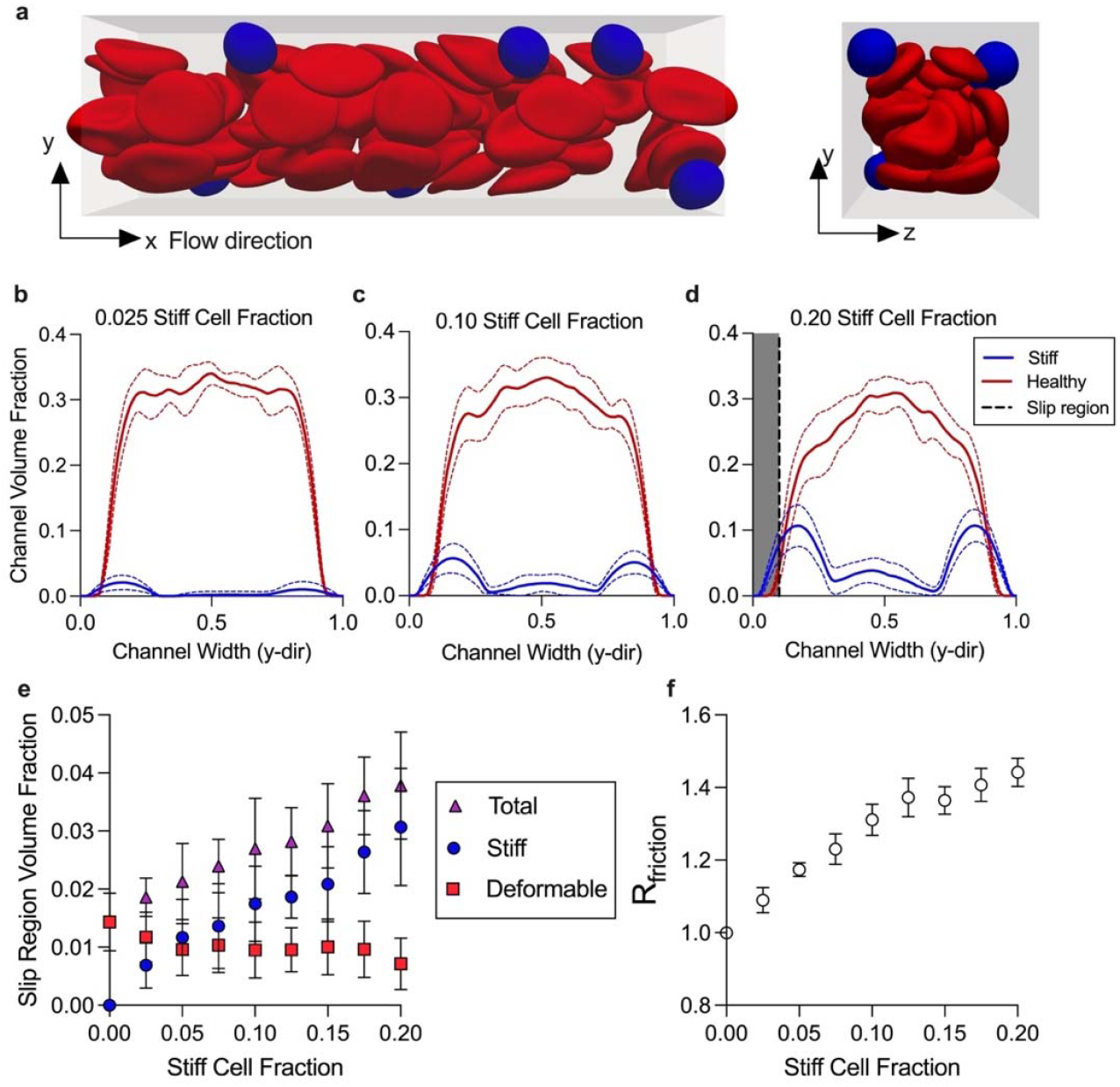
Simulations of heterogeneous blood suspensions in confined flow. a) Simulation data shows spatial localization of the stiff cells (blue) near the channel boundaries compared to the deformable (red) cells. b-d) Local channel volume fractions of the stiff and deformable cells across the channel width for three different stiff cell fractions (0.025, 0.10 and 0.20). Mean (solid line) and 95% confidence (dashed line) plotted for local channel volume fractions of stiff (blue) and deformable (red) cells for n=10 simulations. Due to the finite resolution of the simulations, a slip region (gray), rather than slip length, was defined and held constant. e) The mean volume fraction of stiff (blue circle), deformable (red square) and total (purple triangle) cells within the slip region for each stiff cell fraction is plotted. Error bars representing plus or minus the standard error of the mean (+/-SEM) for n=10 simulations. f) The normalized mean frictional resistance (open circle) plotted against stiff cell fraction. The frictional resistance was calculated using the pressure drop and the slip flow rate, quantified by the average velocity in the slip region. Error bars representing +/-SEM for n=10 simulations.

### In experiments, margination of stiffened RBCs results in rapid localization near channel walls

To test the predictions from the simulations experimentally, we fluorescently stained RBCs from SCD donors and mixed the stained sickle RBCs with unstained RBCs from donors without SCD to a final volume fraction of 25% (Fig. 4a, Methods and Materials 8.0). Upon deoxygenation, only the stained cells become stiff, and the fluorescent signal provides their spatial location. The cells were perfused through a microfluidic device composed of 8 channels, 30 μm wide and 8 μm tall, under a fixed pressure drop of 250 mbar (Fig. 4b). This design provided a quasi-2D configuration in which margination occurs in the imaging plane, and the fluorescent signal is enhanced by eliminating any out of focus effects. Representative images of the fluorescent signals from the stained cells under 21% oxygen and 0% oxygen are shown in Figure 4c and Figure 4f, respectively. The fluorescent streaks represent stained sickle RBCs (streaking is due to the long exposure times needed) (S.I. Video 1 and 2). The spatial distribution of the fluorescent signal under 21% oxygen shows a distribution centered around the middle of the channel suggesting the sickle RBCs concentrate in the middle of the channel when they are deformable (Fig. 4d). However, under deoxygenation, the sickle RBCs concentrate near the boundaries, indicating mechanically induced margination (Fig. 4g). Taken together, these experimental results confirm the predictions from our simulations that margination, due to heterogeneity in mechanical properties, changes the spatial organization of cells and drives the increase in frictional resistance at low stiff cell fractions. Furthermore, we sought to understand the kinetics of margination as it relates to timescales relevant to in-vivo conditions. Therefore, we seeded 25% RBC volume fraction healthy blood with 0.1% volume fraction BSA coated polystyrene beads. We observed rapid margination occurring at the inlet of the microfluidic device. Most beads marginated in the first channel and appeared to concentrate near the channel corners (Fig. 4i, S.I. Video 3). Therefore, we expect margination to occur rapidly in microfluidic flows and contribute to the local rheology measured in sickle blood flow with low stiff cell fractions.

**Figure 4.**
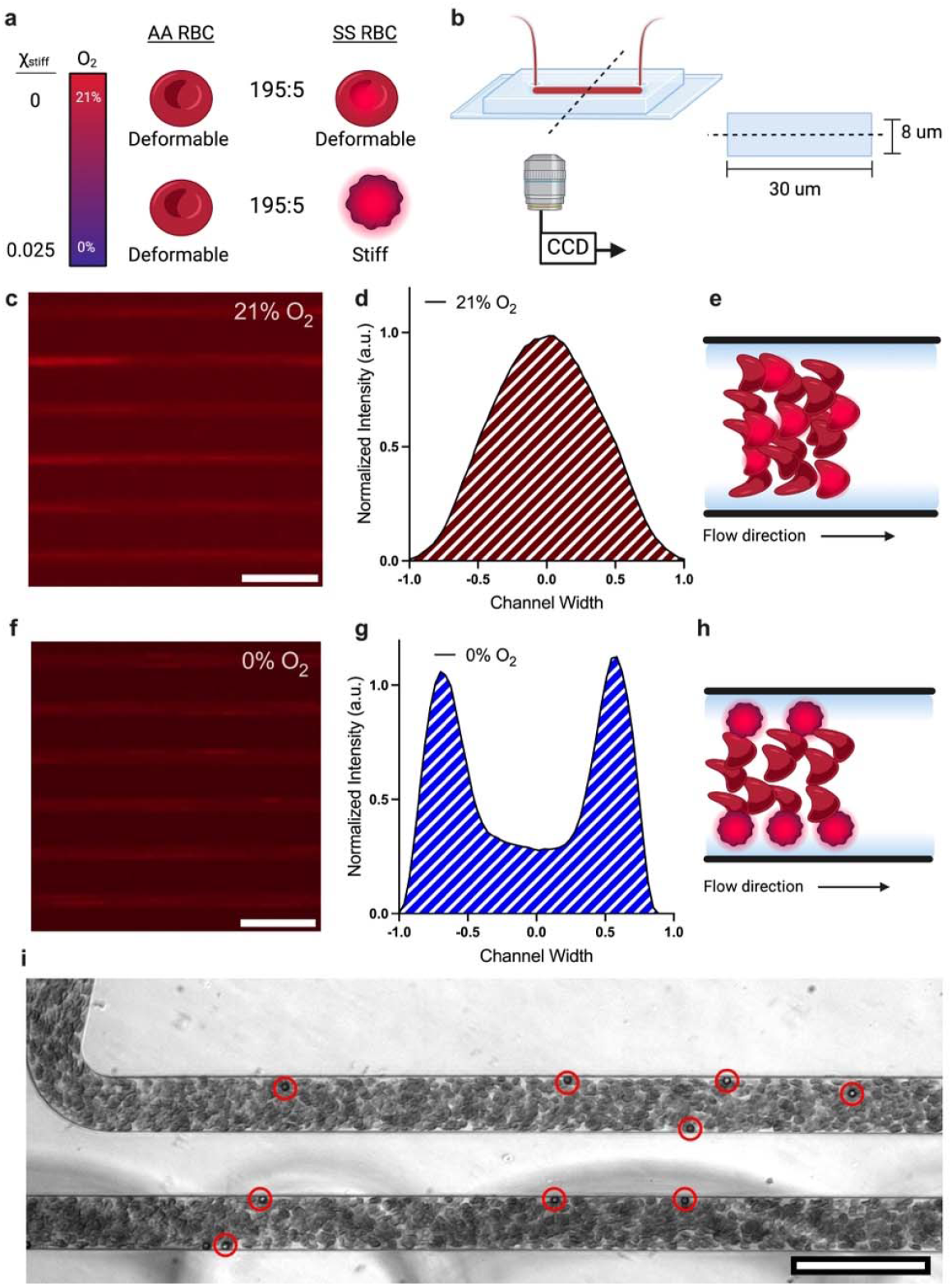
Sickle RBCs marginating in response to hypoxia when present in mixtures of healthy RBCs. a) RBCs from a donor without SCD (AA RBC) are mixed with stained sickle RBCs (SS RBC) at volume ratios of 195:5, respectively. Under 21% oxygen, all cells are deformable. Therefore, the fluorescent signal represents deformable RBC flow. Under 0% oxygen, the sickle RBCs stiffen and provide a fluorescent signal for the spatial distribution of the cells. b) Schematic of the microfluidic channel geometry for blood flow. c) Representative fluorescent images of cells flowing in the microfluidic device under 21% oxygen. Images taken at 5x magnification show 6 of the 12 channels in the device. d) Spatial distribution of fluorescent signal under 21% oxygen for SS RBCs. e) Representative drawing of RBC flow arrangement in heterogeneous mixtures when all cells are deformable. f) Representative fluorescent image of cells flowing in the microfluidic device under 0% oxygen. Images taken at 5x magnification show 6 of the 12 channels in the device. g) Spatial distribution of fluorescent signal under 0% oxygen for SS RBCs. h) Representative drawing of RBC flow arrangement when the SS RBCs are stiff due to hypoxia. i) Experimental image of polystyrene beads marginating in healthy blood. Red circles highlight beads for clarity. Scale bars represent 100 μm. Data shown for a single representative blood sample.

### Axial variations in local hematocrit (or RBC volume fraction) drive extreme profile blunting at high stiff cell fractions

At high values of χ _stiff_, we found the viscous resistance diverges like that of a concentrated suspension approaching jamming (Fig 2d). However, the RBC volume fractions, or hematocrit (HCT), used in our experiments were well below the theoretical jamming volume fraction of 58-64% for rigid particles^36^. To obtain a clearer understanding of the overall blood dynamics in this scenario, we obtained experimental videos of flowing blood over extended length and time scales (Fig. 5a, S.I. Video 4). Interestingly, at low oxygen tensions, variations in the local volume fractions of RBCs along the length of the channel developed, as seen by changes in local light intensity (Fig. 5a). Under fully deoxygenated conditions, we estimate the local HCT corresponding to high and low intensity regions from video data (Methods and Materials 9.0). The results for blood from a representative donor demonstrate that the HCT is no longer uniform along the length of the channel with values as high as 35%, as low as 12%, and a mean HCT of 24% (Fig. 5b). This finding contrasts with the local HCT appearing relatively uniform at 21% oxygen (Fig. 5c). Using the local gradients in the axial HCT signal, we segmented the video data by HCT and measured the local flow features as a function of the local HCT (Fig. 5d). We found the normalized velocity profiles in the high HCT regions to show extreme profile flattening. By contrast, the velocity profiles in the low HCT regions were more parabolic (Fig. 5e). For a representative sample, we found that the profile bluntness, quantified by the wall velocity divided by the maximum velocity (V_wall_/V_max_), clearly depends on the local HCT (Fig. 5f). Meanwhile, the average local velocity (V_avg_) varied to some extent with HCT, however these findings were unsurprising given the spatial-temporal heterogeneity in HCT as well as V_avg_ representing only the average speed of the solid phase (Fig. 5g). In summary, our findings reveal the overall blood flow dynamics measured at high stiff cell fractions are a result of significant axial variations in the local volume fractions of cells which can be observed as temporal fluctuations in volume fraction. Given the extreme blunting in the high HCT regimes, these regions demonstrate an apparent jamming behavior, driving the increases in the macroscopic resistances like that of a concentrated suspension.

**Figure 5.**
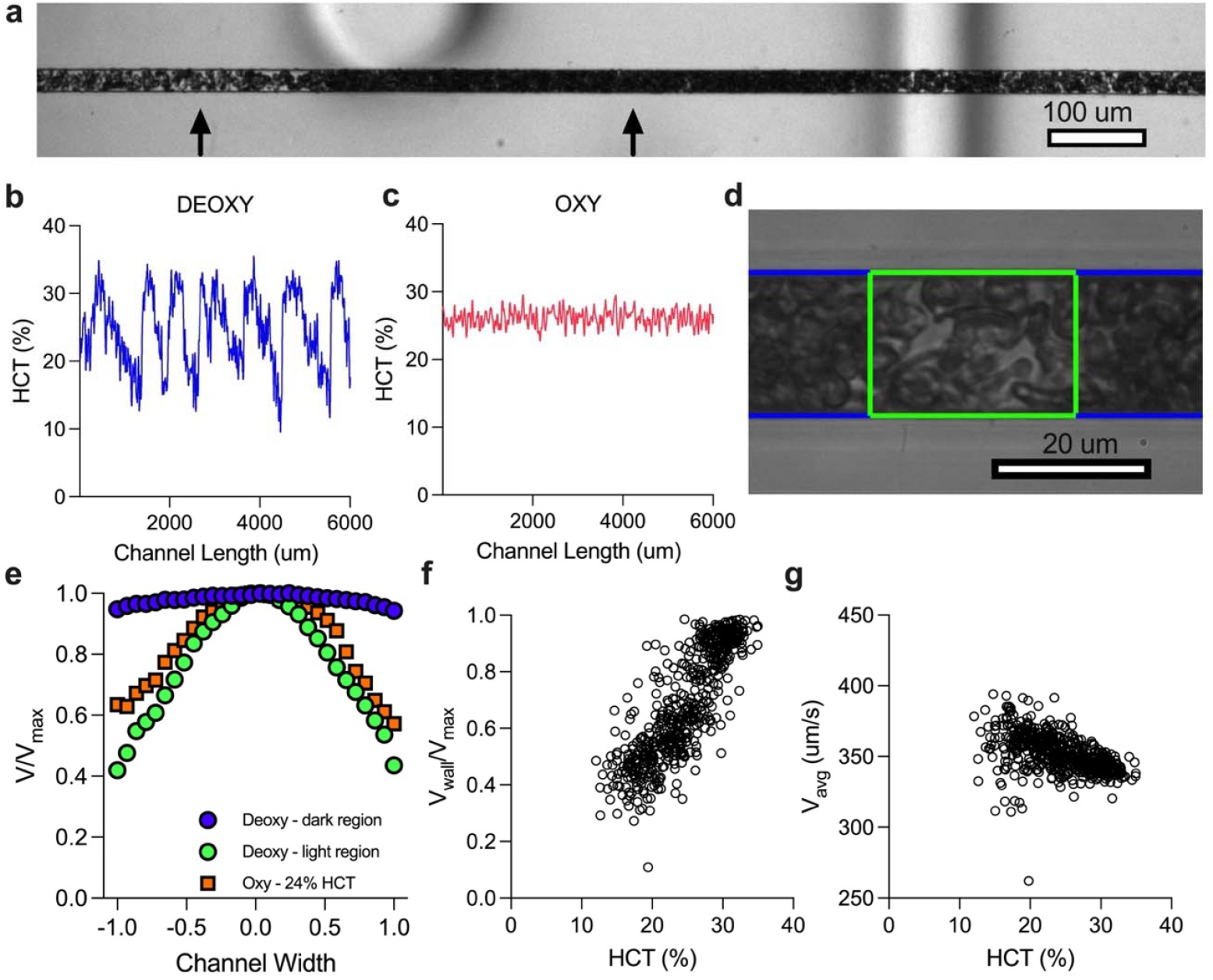
Axial variations in local RBC volume fractions and extreme profile blunting occur at high stiff cell fractions. a) Axial variations in the local RBC volume fractions in the microfluidic channel for sickle blood under 0% oxygen. The bright region has low RBC concentration while the dark region has a higher RBC concentration. Regions are noted by arrows for clarity. b) The local HCT was quantified over the length of the experimental channel. Oscillations in the local HCT fluctuated between 12% and 35%. c) The local HCT across the length of the channel at 21% oxygen. d) Local flow data is chunked into high (blue) and low (green) HCT regions. Local flow profiles within each region are measured and compared to the corresponding HCT. e) Normalized flow profiles from the high HCT region (blue circle) and low HCT region (green circle) are plotted and compared to 21% oxygen (orange square). The normalized flow profiles reveal that extreme blunting occurs in the higher HCT region. f) V_wall_/V_max_ from chunked data is plotted against the corresponding HCT. g) Average speeds from the chunked data plotted against corresponding HCT. Data in panels b,c,e,f, and g were obtained from a single representative SCD blood sample.

## Discussion

Here, we developed a microfluidic platform along with computational modeling to measure and understand how heterogeneity in RBC properties drives blood rheology. Our results show that the extreme variability in blood rheology among individuals with SCD, which is known for heterogeneity in RBC properties, is almost entirely explained by accounting for the distribution of stiff cells in suspension at a given oxygen tension. We find these mechanically heterogeneous mixtures obey rigid-particle suspension mechanics on a macroscopic level. However, the mechanisms driving the rheological behavior are highly dependent on the cross-sectional and axial spatial distribution of cells. Ultimately, these findings reveal an important biophysical mechanism that can explain pathological changes in blood rheology.

### Suspension rheology

Our findings have implications in suspension rheology – blood from patients with SCD is a heterogeneous suspension containing a mixture of stiff and deformable cells, with the relative number of each cell type determined by the oxygen tension in a patient-specific way. Our spatially resolved measurements allow us to extract the overall flow resistance of such suspensions in microchannels, and to decompose the overall resistance into a frictional component, determined by wall slip, and a viscous component, determined by bulk viscosity. We find that all three of these resistances are largely determined by the fraction of stiff cells in the mixed suspension. However, there are significant spatial complexities in suspension organization, which determine how the flow resistances depend on stiff cell fraction. For example, at low stiff cell fractions we see a cross-sectional variance in cell organization – heterogeneity in cell mechanical properties drives margination and localization of stiff cells near channel walls. This leads to a reduction in the cell-free layer thickness, and therefore an increase in frictional resistance, even for very small numbers of stiff cells. This finding contrasts with suspensions of homogeneous rigid particles, where classic theories do not consider frictional effects at very low particle volume fractions in confinement^42,43^. More recent measurements of wall friction in rigid particle suspensions have been performed but only above around 20% volume fraction of particles^33^. Our results concerning frictional resistance demonstrate the combined importance of particle organization and geometric confinement in determining the behavior of heterogeneous particle suspensions.

At higher stiff cell fractions, we observe significant axial heterogeneity in the volume fraction of cells. These heterogeneities generate regions of high and low volume fraction, with different flow profiles. We observe extreme bluntness of the profiles in the high-volume fraction regions, which qualitatively reflects an apparent jamming behavior in the bulk portion of the suspension^47^. Despite the local packing, we find that the volume fractions in these regions are significantly lower than classical measurements of the jamming fraction for suspensions of rigid particles^34,36^. However, we note that previous measurements are typically performed on frictionless, unconfined particle suspensions. We suspect additional effects such as polydispersity, confinement, or cell-cell friction may contribute to the extreme bluntness. In particular, recent theory and simulations have found that particle roughness can generate significant extra effective friction^48–50^. Taken together, our findings highlight the importance of spatial particle organization in determining the rheology of suspensions flowing through confined geometries.

### Pathophysiology

Here, we demonstrate hypoxia-induced margination of unaltered RBCs from SCD donors. Given we detect stiff cells at physiological oxygen tensions as high as 12% oxygen, our results suggest that small amounts of these stiff cells are likely present throughout the vasculature, and that margination of these cells occurs even at low abundance. This corroborates the findings of previous work that stiffened RBC margination in SCD is an important biophysical contribution to systemic vascular inflammation and disease pathology^51^. Furthermore, we observe that the process of margination of stiff RBCs increases friction and counteracts the Fahreaus effect, thereby increasing the effective viscosity by increasing the concentration of cells near channel walls in the microvasculature. We postulate that small amounts of stiffened cells may have a significant effect on peripheral vascular resistance and their presence near blood vessel walls may alter mechanical homeostasis in the vasculature^52^. Further work should be aimed at understanding the dynamics of these small populations of stiff cells in more complex vascular networks to provide insight into their impact on network level flow dynamics, possibly indicating which vascular structures might be most susceptible to margination induced inflammation, and to help guide more complex in-vivo studies^53^. The physiologic implications of our results demonstrating axial-temporal variation in HCT under extreme hypoxia is less clear. However, we note that such axial variation has been observed in blood flow imaging in the vasculature of transgenic sickle mouse models^54^. Thus, we conclude that this is not an experimental artifact but a real phenomenon, and more work is warranted to understand the potential clinical impact.

### Clinical Relevance and Outlook

Here, we show that the hypoxia-driven rheological responses across multiple blood donors appear remarkably similar when accounting for the distribution of cells in the polymerized hemoglobin S state. Our results also demonstrate that these results can account for medical interventions such as exchange transfusion therapy, which dramatically alter the fraction of stiffened RBCs. Additionally, therapies such as hydroxyurea or gene therapy, which increase fetal hemoglobin as a therapeutic strategy, also modulate patient specific stiff RBC distributions^40^. Therefore, our findings establish a strong link between therapies, which often have variable effects on individuals, and measurable aspects of the blood, that can account for variability between individuals based on their response to therapy. These findings further motivate the need for single RBC measurements when assessing therapeutic strategies for individuals with SCD^55^. Beyond SCD, diseases such as malaria and diabetes also compromise the mechanical deformability of the RBC, and mechanisms driving increases in blood viscosity are attributed to increases in RBC stiffness^56,57^. Evaluating pathological rheology as it relates to heterogeneity in the properties of RBCs is likely relevant for a range of diseases, and the results here provide a framework to understand how pathologies may evolve across those diseases.

## MATERIALS AND METHODS

### 1.0 Blood sample collection and preparation

Blood samples from healthy donors and donors with sickle cell disease (SCD) were collected at the Massachusetts General Hospital under Institutional Review Board approved protocols (2006P000066), at the University of Minnesota and Children’s Hospital and Clinics of Minnesota under Institutional Review Board approved protocols (STUDY00003). All human subjects gave informed consent before participating in this study. Complete blood counts (CBC) and hemoglobin variant fractions HbA, HbS, HbA2 and HbF are included in S.I. Table 1.

### 2.0 Blood preparation

All blood samples used for experiments were collected in 0.109M sodium citrate buffer (BD cat. no. 363083) and stored up to four days at 4 °C prior to testing. For rheological measurements, blood plasma was removed using a three-step wash and centrifugation procedure at 400 x g. Washed red blood cells were resuspended in 2% bovine serum albumin (Sigma Aldrich cat. no. A7030-50G) in Dulbecco’s phosphate buffer saline (PBS) (Corning cat. no. 21-031-CV) to a target of 25% hematocrit (HCT) or RBC volume fraction. The HCT target was chosen based on the average hematocrit for SCD patients. For single cell measurements, 12 μL of packed, washed RBCs were added to 288 μL of 25% HSA solution (Gemini Bio cat. no. 800-120) and 12 μL of 0.8 g/dL acid blue 9 in 1x PBS (AB9, TCI America, Portland, OR, cat. no. B0790). For exchange transfusions, washed healthy donor blood (HbA only blood) was suspended to a target HCT of 25% in 2% BSA and mixed at different volume ratios with 25% HCT HbS blood. Samples chosen for transfusion were type matched and had similar MCV values. Packed RBC subsamples of these mixtures were used for single cell measurements.

### 3.0 Microfluidic device manufacturing

All experimental studies were conducted using polydimethylsiloxane (PDMS) devices. Briefly, the microfluidic rheology device consists of three layers bonded together. The first layer is composed of two gas chambers 150 μm tall allowing for independent control of oxygen to either side of the device. The second layer is a 100 μm tall hydration layer. The third layer is a composed of a 40×20 μm resistor that bifurcates into two 20×20 μm channels, 22 mm in length. The single-cell microfluidic device was manufactured using the exact design from Di Caprio et al^39^. The margination microfluidic device design was adapted from the design of Di Caprio et al. using a blood layer composed of 21 channels each 8 μm tall, 30 μm wide and 40 mm long. Silicon wafer molds were fabricated using negative resist photolithography for the indicated feature geometries. Each PDMS layer was fabricated by soft lithography from silicon wafer molds. PDMS elastomer and curing agent were mixed using a 10:1 ratio by weight (Sylgard 184, Dow Corning, USA). Microfluidic layers were cured at 75°C for 2 hours, bonded on the same day, and used within 24 hours of manufacturing. Each layer was plasma bonded at 10 cc/min air flow rate, 75% power, and 60 seconds exposure time settings (PE-50, PlasmaEtch) and cured at 120°C for 5 minutes. The merged layers were then plasma bonded to a clean glass microscope slide using the same plasma settings and cured at 120°C for 5 minutes.

### 4.0 Microfluidic rheology platform, measurement system and flow rate validation

Using a microfluidic platform previously describe, flow measurements were made in a 20×20 μm channel under varying degrees of hypoxia^41,58^.To characterize each blood sample, the microfluidic rheology device was visualized on a Zeiss Axio Vert microscope using a 40x objective (Zeiss 40x/0.6NA, air) encased in a 37°C environmental incubator chamber. Blood was perfused at a fixed pressure drop and initial average velocity of 900 μm/s Images were acquired using a high-speed camera (Phantom Miro C-110) at 400 fps for a 512×1280 image at a region of interest near the end of the experimental channel. Oxygen (21% O2, balance N2) was cycled from 21% to the desired hypoxic condition by mixing with pure nitrogen using a solenoid valve gas mixer reported previously^58,59^. Blood rheology was measured by directly visualizing the blood flow and calculating local velocity profiles using computer vision techniques. Briefly, the total RBC flow rate, *Q*_*total*_, in the experimental channel was calculated assuming the velocity profile corresponds to a height-averaged velocity profile across the device therefore, *Q*_*total*_ *= V*_*avg* ×_ *width* × *height*. During 21% oxygen conditions, the experimental and bypass channel are assumed to have the same flow rate. An external flow meter (Flow Unit XS, Fluigent) was then incorporated to validate flow rates measured using the computer vision techniques. We found no significant difference between the flow meter reading and computer vision estimates at different time points corresponding to 21% oxygen during our experiments, when both channels have the same flow rate (S.I. Figure 1). Effective, friction and viscous resistances were calculated from the flow rates using methods described previously^41^. Briefly, *R*_*friction*_ *=* Δ*P/Q*_*slip*_ and *R*_*viscous*_ *=* Δ*P/Q*_*bulk*_. Δ*P* is the pressure drop across the experimental channel, *Q*_*slip*_ is the slip flow rate, and *Q*_*bulk*_ is the bulk flow rate. *Q*_*slip*_ is calculated from the wall velocity multiplied by the channel cross-sectional area. *Q*_*bulk*_ is simply the difference of *Q*_*total*_ and *Q*_*slip*_. Here, we normalized the resistance calculations to the corresponding 21% oxygen value immediately preceding the deoxygenation cycle step so that any effects due to temporal drift were minimized.

### 5.0 Single-cell microfluidic platform

Red blood cells containing HbS polymer at various oxygen tensions were identified using a high-throughput single cell microfluidic platform following the methods previously described^39^. Experiments were run on the same day as the blood rheology experiments. Blood was prepared as described in Materials and Methods 2.0. Prepared blood was visualized using a Zeiss Axio Vert microscope with a 40x/0.75NA air objective encased in a 37°C environmental incubator chamber. Images of single red blood cell at 3 different wavelengths are captured on a color camera (Flir Blackfly S). 610 nm was used to measure RBC volume by dye exclusion and the 405nm/430nm were used to quantify the amount of oxy and deoxy hemoglobin based on the relative absorbance at each wavelength. The RGB images were then classified as either stiff (containing HbS polymer) or deformable (no polymer) using a Resnet50 algorithm previously validated and tested^40^. Population distributions of at least 1000 cells were used to estimate polymerized cell fractions ranging from 0 to 1 for each oxygen tension.

### 6.0 Data analysis, statistics and reproducibility

Data were fit to a n-order polynomial using the least squares regression method. The order of polynomial was chosen as the lowest value in which the r-square value was minimal without over-fitting the data and maintaining a monotonic form. The curve fit (solid lines) and 95% confidence intervals (dashed lines) were plotted. The sum of squared errors (SSE) was then used to compute the variance, defined as the SSE/(n-1), and compare between plots. Statistical analysis was performed in Prism v9.5.0. All experimental measurements used in this study were taken from different blood samples. For simulations, 10 simulations were run for each condition. Error bars representing plus or minus the standard error of the mean (+/-SEM) for n=10 simulations. Mean (solid line) and 95% confidence (dashed line) are plotted for local channel volume fraction estimates. Flow data were analyzed in MATLAB 2022b. Single-cell data were analyzed in MATLAB 2021a and Python 3.10.11.

### 7.0 Red blood cell simulation methods

We performed simulations using a previously developed numerical model: a hybrid immersed-boundary lattice-Boltzmann finite-element model^60^. The suspending fluid was solved by a lattice Boltzmann solver for a geometry of 64 × 20 × 20 μm ^3^ (288 × 90 × 90 grid points), periodic in the x-direction, with a viscosity of 0.001 Pa s, and an imposed pressure gradient of 1.3 × 10^5^ N/m^3^ (c.f. an experimental pressure gradient of 1.44 × 10^5^ N/m^*3*^). Healthy red blood cells were represented by a finite-element triangular mesh with 6480 facets, with equilibrium shape as a biconcave disk of radius 18 grid points (4 μm). The membrane model of the RBC was as in <u>Krüger</u> et al.^45^, parameterized by (in simulation units) (κ_S_, κ_B_, κ α, κ_A_, κ_V_) = (0.00132, 0.00107, 0.75, 1, 0.75), corresponding to healthy RBC with shear modulus 5 × 10^−6^ N/m. Stiff red blood cells were represented by a triangular mesh with 2880 facets, with equilibrium shape as a sphere of radius 12 grid points (∼2.67 μm enclosed volume is approximately 80% of healthy cell enclosed volume), parameterized by (κ_S_, κ_B_, κα, κ_A_, κ_V_) = (0.0132, 0.00474, 0.75, 1, 0.75), corresponding to a shear modulus ten times greater than the healthy RBC shear modulus (5 × 10^*-5*^ N/m). In our simulations we fixed the cell volume fraction as 25%, resulting in simulations run with the number of stiff:healthy cells as (0:63, 2:61, 4:60, 6:58, 8:57, 10:55, 12:53, 14:52, 16:50). To improve numerical stability and avoid particle overlap, a short-range repulsive force between cells was used. This force was zero for distances larger than 2 grid points and behaved as 1/*r*^2^ for shorter distances. To improve statistics, all simulations were repeated ten times, with random initial RBC configurations. Statistics were calculated after the simulation had been run for 1.0 × 10^6^ time steps (equivalent to 0.5 seconds), to ensure that a quasi-steady state had been attained (verified by confirming convergence of calculated statistics). Cell volume distributions were calculated by finding the enclosed cell volumes bounded by parallel planes located at y=0 and y=Y, for Y in [0,90], at timestep 10^6^. Wall frictional resistance was defined as 1/*V*_*Wall*_, normalized by the frictional resistance for the zero stiff cell case (100% deformable cells). V_*Wall*_ is the wall velocity, defined as the mean x-velocity of the nodes with y in grid point boundaries [0,9) or (81,90] (within 10% channel width of the wall), for 10^6^< t < 1.6 × 10^6^.

### 8.0 Fluorescent cell staining protocol and analysis

RBCs from an un-transfused SCD blood sample was fluorescently stained by pipetting 5 μL whole blood from sodium citrate tubes into 995 μL of a 1/1000 dilution of CellMask DeepRed (C10046) in 1x PBS. The blood stain mixture was incubated for 5 minutes, spun at 400 x g for 3 minutes, and the supernatant was removed. Stained RBCs were resuspended in 1 mL PBS and then spun at 400 x g for 3 min. The supernatant was removed, and the remaining pellet was resuspended with 195 μL of 25% HCT healthy RBC in 2% BSA in PBS suspension yielding approximately a 0.025 fraction of HbS red blood cells in 25% HCT blood suspension. Blood was perfused in a 2-layer microfluidic device consisting of a blood layer with 12 straight channels 30 μm wide by 8 μm tall and a gas layer. Fluorescent images were acquired using a Zeiss Axio Vert microscope enclosed in a 37°C environmental incubator chamber using 20x and 5x objectives (Zeiss APO 20x/0.8NA and Zeiss 5x/0.16NA objective) and imaged using a CMOS camera (ORCA-Flash4.0LT, Hamamatsu). 300 frames at 21% oxygen and 0% oxygen were captured for each oxygen tension. Image fluorescent intensity data was processed by averaging the lateral (30 μm width direction) fluorescent intensity signal across the axial length (flow direction) of the channel using ImageJ to enhance contrast and remove background and custom scripts in MATLAB 2022b for quantification of the processed image stacks. Intensity plots in Figure 3d and 3g were each normalized by their respective area under the curve.

### 9.0 Hematocrit estimation protocol and validation

We estimated the total volume fraction of RBCs in the microfluidic channel using a method described by Roman et al., whereby the optical density of absorbed light of red blood cells is related to the local HCT by counting the cells and multiplying by the RBC volume in the given channel volume^61^ (S.I. Fig. 3).Two standard curves were measured for 21% and 0% oxygen given the differences in absorbance for oxy and deoxy hemoglobin (S.I. Fig.3). Similar to Roman et al., we successfully counted cells up to a HCT of 12%. In attempt to validate and test the dynamic range of our measurement, we flowed packed red blood cells into the device. The estimated feed HCT from a sample of the inlet tubing was 57% measured by hemacytometer. The tube HCT was estimated using the calibration curve generated in S.I. Fig. 3a to be 41%. However, due to the known Fahraeus effect, we expect the tube HCT to be lower than the feed HCT for a 20×20 μm channel. Therefore, we used the relationship proposed by Pries et al.^26^ which relates the feed HCT to tube HCT and estimated the tube HCT to be 45% given a feed of 57%, giving us reasonable confidence to the validity and dynamic range of our measurements.

## Materials and Data Availability

All experimental protocols are described in the Materials and Methods. The authors declare that the data supporting the findings of this study are available within the paper and its Supplementary Information files. Data associated with patient specimens that can be shared under our IRB Protocol are included in the Supplementary Information (S.I. Table 1). Should any raw data files be needed in another format they are available from the corresponding author upon reasonable request.

## Code Availability

Blood flow analysis code is available at https://doi.org/10.5281/zenodo.14934042. Simulation software and single-cell code is as described in Krüger et al. and Di Caprio et al., respectively^39,45^.

## Acknowledgments

This work was supported by the National Heart, Lung, and Blood Institute Grants HL130818, HL132906, T32-HL139341 and the National Science Foundation Graduate Research Fellowship Program under Grant No. 2237827. Portions of this work were conducted in the Minnesota Nano Center, which is supported by the National Science Foundation through the National Nano Coordinated Infrastructure Network under Award ECCS-1542202. Portions of this work were supported by a research grant from the UKRI Engineering and Physical Sciences Research Council (EP/T008806/1). We thank our clinical teams for assistance with blood sample collection, specifically, Chhaya Patel and Simran Singh from Massachusetts General Hospital, Ali Kolste and Emma Smith from Children’s Hospital and Clinics of Minnesota, and Yvonne Data M.D. from the University of Minnesota. PP was supported by a UKRI Future Leaders Fellowship (MR/V022385/1). The authors acknowledge the use of the UCL Myriad High Performance Computing Facility (Myriad@UCL), and associated support services, in the completion of this work.

## Notes

### Competing Interest Statement

The authors have declared no competing interest.

