## Supplementary Information for "Suspension physics govern the multiscale dynamics of blood flow in sickle cell disease"

**S.I. Video 1** Video of spatial distribution of fluorescent signal under 21% oxygen for fluorescent SS RBCs. Videos acquired at 20 fps and 20x magnification. Playback speed at 20 fps.

**S.I. Video 2** Video of spatial distribution of fluorescent signal under 0% oxygen for fluorescent SS RBCs. Videos acquired at 20 fps and 20x magnification. Playback speed at 20 fps.

**S.I. Video 3** Video of actively marginating polystyrene beads in 25% hematocrit blood at 21% oxygen. Video acquired at 400 fps and 10x magnification. Playback speed at 40 fps.

**S.I. Video 4** Video of axial variations in hematocrit at 0% oxygen. Video acquired at 400 fps and 5x magnification. Playback speed at 40 fps.

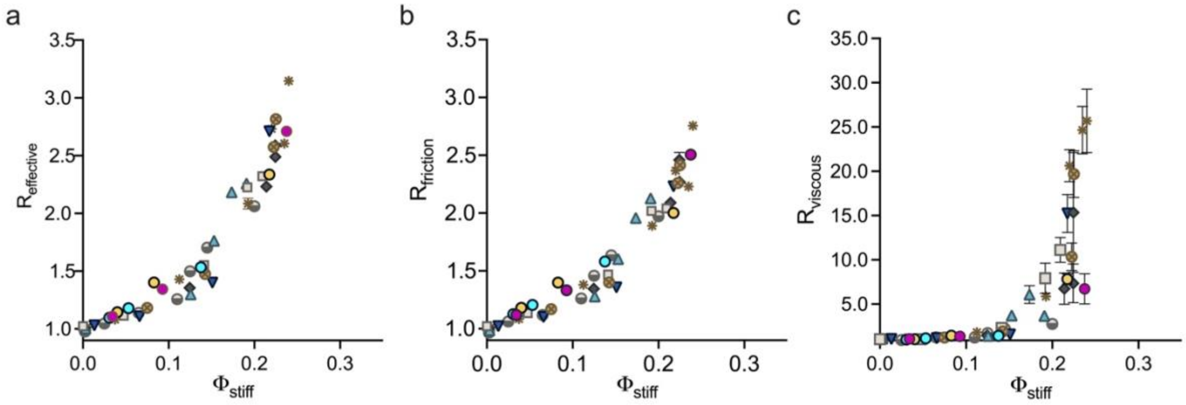


**S.I. Figure 1** Effective rheology scaled to volume fraction of stiff RBCs, $\Phi_{stiff}$, where the total volume fraction of RBCs (stiff + deformable) is fixed, $\Phi = 0.25$. a) $R_{effective}$ plotted as a function of $\Phi_{stiff}$. b) $R_{friction}$ plotted as a function of $\Phi_{stiff}$. c) $R_{viscous}$ plotted as a function of $\Phi_{stiff}$. Results shown for n=10 blood samples from SCD donors. Error bars represent plus or minus the standard error of the mean (+/- SEM) for resistance data gathered for n = 11 independent sampling time points during flow data acquisition. Error bars smaller than the data symbol are not shown.

**S.I. Table 1** Patient hematological data. Patient genotype, hemoglobin fractions and complete blood count data are included for n=10 blood samples from SCD donors. Acronyms: Hemoglobin S (HbS), hemoglobin A (HbA), hemoglobin F (HbF), hemoglobin A2 (HbA2), white blood cell count (WBC), mean corpuscular volume (MCV), mean corpuscular hemoglobin concentration (MCHC), and red blood cell distribution width (RDW).

| Patient ID | Genotype | HbS (%) | HbA (%) | HbF (%) | HbA2 (%) | WBC (K/uL) | Hemoglobin (Hgb) (g/dL) | Hematocrit (Hct) (%) | MCV (fL) | MCHC (%) | RDW (%) |
|---|---|---|---|---|---|---|---|---|---|---|---|
| Sample1 | HbSS | 80.1 | 0 | 16.7 | 3.2 | 14.6 | 8.3 | 24.2 | 91 | 34.3 | 16.6 |
| Sample2 | HbSS | 58.5 | 0 | 40.1 | 1.4 | 4.2 | 9.5 | 25.1 | 113 | 37.8 | 15.3 |
| Sample3 | HbSS | 80 | 0 | 16.5 | 3.5 | 6.6 | 7.7 | 22.5 | 69 | 34.2 | 16 |
| Sample4 | HbSS | 90.6 | 0 | 5.8 | 3.6 | 9 | 8.3 | 23.7 | 98 | 35 | 22 |
| Sample5 | HbSS | 75.7 | 18.5 | 2.8 | 3 | 10.4 | 7 | 19.3 | 87 | 36.3 | 23.8 |
| Sample6 | HbSS | 67.9 | 0 | 29.1 | 3 | 6.2 | 9.6 | 28.9 | 91 | 33.2 | 19.1 |
| Sample7 | HbSS | 89.9 | 0 | 7.2 | 2.9 | 8.1 | 10.3 | 28.9 | 76 | 35.6 | 18.6 |
| Sample8 | HbSS | 50.6 | 35.4 | 11.2 | 2.8 | 8.95 | n.d. | 27.3 | 97.5 | 34.8 | n.d. |
| Sample9 | HbSS | 89.2 | 0 | 7.4 | 3.4 | 11.1 | 9.8 | 27.8 | 96 | 35.3 | 17.8 |
| Sample10 | HbSS | n.d. | n.d. | n.d. | n.d. | 5 | 10 | 29.5 | 101 | 33.9 | 17.8 |

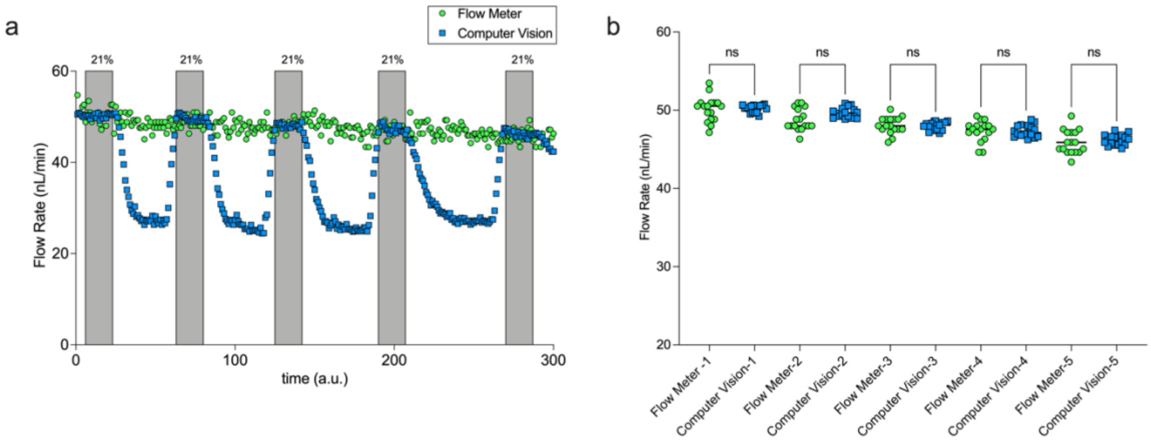


**S.I. Figure 2** Flow rate validation between flow meter and computer vision flow rate estimates. a) Temporal data showing flow rate readings from the Fluigent flowmeter (green circle) and flow rate estimates made using the average velocity obtained from computer vision estimates multiplied by the cross-sectional area (blue square). b) Flow rate readings show good agreement when blood is exposed to 21% oxygen as the are no significant differences between the flow meter and computer visions. An ordinary one-way ANOVA with multiple comparisons was performed in Prism v9.5.0 where $p<0.05$ indicates significance.

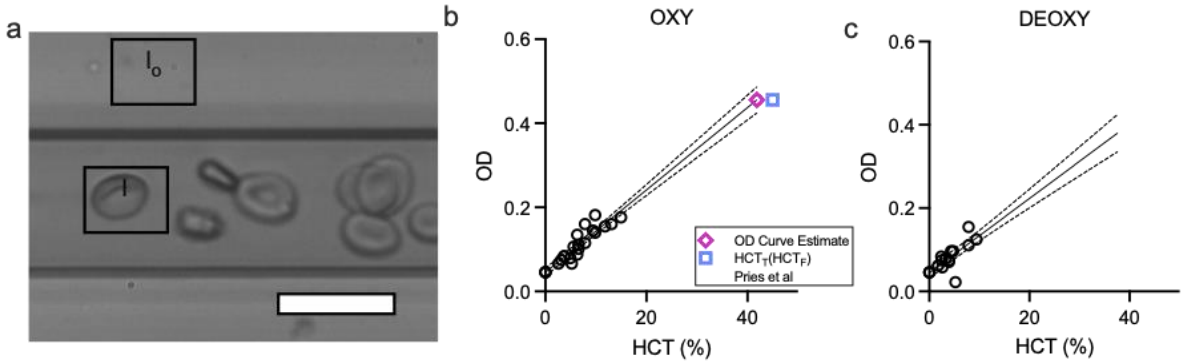


**S.I. Figure 3** Hematocrit estimation and validation. a) The optical density of absorbed light of red blood cells is related to the local HCT by counting the cells and multiplying by the mean RBC volume (the mean corpuscular volume is obtained from complete blood count data) in the given channel volume. I is the absorbed light. $I_o$ is the incident light. The channel volume is calculated using the area of the region of interest (black box) multiplied by the channel depth (20 um). b) Optical density (OD) plotted against estimated hematocrit (HCT (%)) for RBCs under 21% oxygen. Tube HCT of a dense RBC suspension estimated from the calibration curve (pink diamond). Tube HCT predicted from the feed HCT (blue square) by adjusting for the difference in tube and feed HCT for small channels. c) OD plotted against estimated HCT (%) for RBCs under 0% oxygen.
